# Biologging of emperor penguins – attachment techniques and associated deployment performance

**DOI:** 10.1101/2021.06.08.446548

**Authors:** Aymeric Houstin, Daniel P. Zitterbart, Alexander Winterl, Sebastian Richter, Víctor Planas-Bielsa, Damien Chevallier, André Ancel, Jérôme Fournier, Ben Fabry, Céline Le Bohec

## Abstract

An increasing number of marine animals are equipped with biologgers, to study their physiology, behaviour and ecology, often for conservation purposes. To minimise the impacts of biologgers on the animals’ welfare, the *Refinement* principle from the Three Rs framework (*Replacement, Reduction, Refinement*) urges to continuously test and evaluate new and updated biologging protocols.

Here, we propose alternative and promising techniques for emperor penguin (*Aptenodytes forsteri*) capture and on-site logger deployment that aim to mitigate the potential negative impacts of logger deployment on these birds. We equipped adult emperor penguins for short-term (GPS, Time-Depth Recorder (TDR)) and long-term (*i*.*e*. planned for one year) deployments (ARGOS platforms, TDR), as well as juvenile emperor penguins for long-term deployments (ARGOS platforms) in the Weddell Sea area where they had not yet been studied.

We describe and qualitatively evaluate our protocols for the attachment of biologgers on-site at the colony, the capture of the animals and the recovery of the devices after deployment. We report unprecedented recaptures of long-term equipped adult emperor penguins (50% of equipped individuals recaptured after 290 days). Our data demonstrate that the traditional technique of long-term attachment by gluing the biologgers directly to the back feathers is detrimental to the birds. It causes excessive feather breakage and the loss of the devices at an early stage. We therefore propose an alternative method of attachment for back-mounted devices. This technique led to successful year-round deployments on 37.5% of the equipped juveniles. Finally, we also disclose the first deployments of leg-bracelet mounted TDRs on emperor penguins.

Our findings highlight the importance of monitoring potential impacts of biologger deployments on the animals and the need to remain critical towards established and new protocols.

## Introduction

Over the last decades, biologging technology -the “use of miniaturised animal-equipped tags for logging and/or relaying data about an animal’s movements, behaviour, physiology and/or environment” ^1^ -has rapidly progressed and led to fundamental advances in ecology of *e*.*g*. terrestrial ^2–4^ and marine predators ^5–7^ including seabirds ^8–17^. This technical evolution that included miniaturisation, design optimisation, storage capacity and power consumption, was supported by the development of new analytical techniques and processing software ^18,19^.

Biologgers can cause discomfort to the tagged animal and may even impede their movements, especially in the case of diving seabirds like penguins where the increased water drag can increase the energy expenditure ^20–24^. However, the miniaturisation of devices ^25^, the establishment of guidelines ^26,27^ and the activities of study review boards that oversee the ethical treatment of animals in scientific studies ^28–31^ help to mitigate negative impacts and to comply with the Three Rs framework (*Replacement, Reduction, Refinement*) ^32^.

Yet, especially in the case of penguin tracking studies, the inability to observe the animals carrying the devices at sea bears the risk that deleterious effects may not be obvious ^23^ or may even remain unnoticed if birds are not resighted. For instance, after decades of flipper banding thousands of penguins ^33^ (and see references in Jackson et al. ^34^), it was only in the 2000’s that studies ^35–38^ assessed its long-term effect, and showed that flipper bands dramatically decreased the survival and breeding success of their carrier. This finding raised important questions about ethics and bias in scientific studies; an issue already highlighted by Wilson et al. ^39^ in 1986. Flipper banding of penguins is a prime example of why it is necessary to study potential impacts of device deployments on animals. It has to be noted that of the five studies ^40–44^ where emperor penguins (*Aptenodytes forsteri*) have been tagged with external biologgers for a year-round deployment duration, none has reported the recovery of the device or a sighting of an equipped bird after deployment. The causes of signal loss remained unclear ^42,45^ and the fate of the device-carrying birds uncertain.

Nonetheless, data obtained from biologgers are often of such importance for conservation biology that the benefits may outweigh the risk for the animals ^46,47^, if the risks to animals are kept minimal. For example, tracking studies that determine the home range and movement corridors of species are often a prerequisite for conservation management policies ^48–51^ as demonstrated by the establishment of the Ross Sea Marine Protected Area (MPA) in 2017. This first MPA adjacent to Antarctica was partly justified by the range of Adélie penguins (*Pygoscelis adeliae*) during their energy-intensive premoult period ^52,53^.

Emperor penguins have not yet been tracked in the Weddell Sea and in the Atlantic sector of the Southern Ocean, thus not much is known about their distribution in these areas. To improve the scientific knowledge about this species and to provide data in support of the development of a MPA in the Weddell Sea area, we have equipped adult and juvenile emperor penguins. Biologger types were chosen according to our research questions and subject to seasonal constraints. We document for the first time the resighting and recapture of long-term equipped emperor penguins as well as device retrieval. Indeed, while back-mounted loggers have already been successfully used for long-term deployments on emperor penguins ^43,44^, their physical impact has never been assessed presumably due to the logistical difficulties in resighting the birds before the annual moult.

Furthermore, we present the first leg-band biologger attachment and deployment on emperor penguins. Several leg-band devices had been successfully tested and deployed on other penguin species (Adélie and macaroni (*Eudyptes chrysolophus*) penguins, see ^11,30,54,55^) and it was shown that the leg-band devices minimised drag, induced little behavioural disturbance and did not jeopardize birds’ survival. To date, no such deployment had been reported on emperor penguins. Additionally, we describe and discuss methods for catching, handling or retrieving (resight and recapture) emperor penguins. These necessary procedures lack standardisation across studies. Some use a rugby-like catch method ^56,57^, others would use a crook ^42,58^ or a fixed enclosure ^59,60^, and the impacts of these procedures on the targeted bird are rarely reported. Summarising, in this manuscript, we describe and review protocols for on-site capture, handling and release of emperor penguins, biologger attachment and recovery techniques that aims to minimise the impacts on the birds’ welfare.

## Methods

### Study site, species, and deployments

This study was conducted at the Atka Bay emperor penguin colony (70°37’S, 08°09’W) in close vicinity (∼ 10 km) of the German research base Neumayer Station III (70°39’S, 08°15’W) during two consecutive summer campaigns (November to January 2017-2019). During these campaigns, we deployed biologgers for short-and long-term deployments. Monitoring periods of weeks to months in summer ^61–64^ are referred to as “short-term”, while year-round planned monitoring that include austral winter are referred to as “long-term” ^62,65,66^.

The deployment protocols possible to implement on emperor penguins largely depend on the species’ phenology (and logistic constraints). The Emperor penguin is the only bird species breeding during the austral winter ^67^, almost exclusively on sea ice ^68^ all around Antarctica ^69^. After a courtship period in March and April, depending on the colony’s latitude, and an incubation period of around 64 days, the chicks hatch in the middle of the austral winter. As central place foragers ^70^, male and female do alternate trips at sea to find food for their sole offspring. By October, the chick is thermally independent and is left on its own while both parents go foraging at sea and return to feed their chick independently ^67,71^. These recurring returns of each adult to feed its chick, approximately once per week, allow deployment and retrieval of short-term data loggers. In December or January, chicks moult and fledge. By the end of the austral summer, the adult emperor penguins moult. For both, moulted chicks (*i*.*e*. juveniles) and adults, a reliable attachment of long-term logging devices on their back is only possible after moulting is largely completed. The majority of juvenile birds will not return to the colony for at least two years ^72^ and previous studies suggest that most of the adults moult on the pack ice ^62,73–75^. There is also no certainty that adults moulting at the colony are actual breeders from that particular colony and that they will return in the next season. Therefore, successfully retrieving the devices is unlikely and the use of transmitting devices is by far the most prevalent technique to ensure data collection.

In this study, we used two capture methods (corral or crook, see the capture protocols section for details) to catch three categories of birds (a pair of an adult with its chick in November/December, juveniles and moulted adults in January) in order to deploy and/or recover six different types of loggers (see Additional file 1). Depending on the duration (short-or long-term) of planned equipment, biologgers were attached by one of four techniques (back-attachment-tape/-cyanoacrylate-glue/-tape-epoxy and leg-band). The four deployment protocols are briefly presented below and summarised in Table 1. Additionally, all birds, *i*.*e*. adults and chicks, were marked with subcutaneous passive integrated transponder (PIT of 3.85 × 32 mm and 0.8 g, Texas Instruments Remote Identification System, TIRIS, Texas, USA) implanted between the tail and left leg (Additional file 2) allowing remote identification of individuals with automatic reading systems. All protocols adhered to current best-practise standards to reduce the risk of physical harm and stress to individuals and the colony.

**Table 1.**
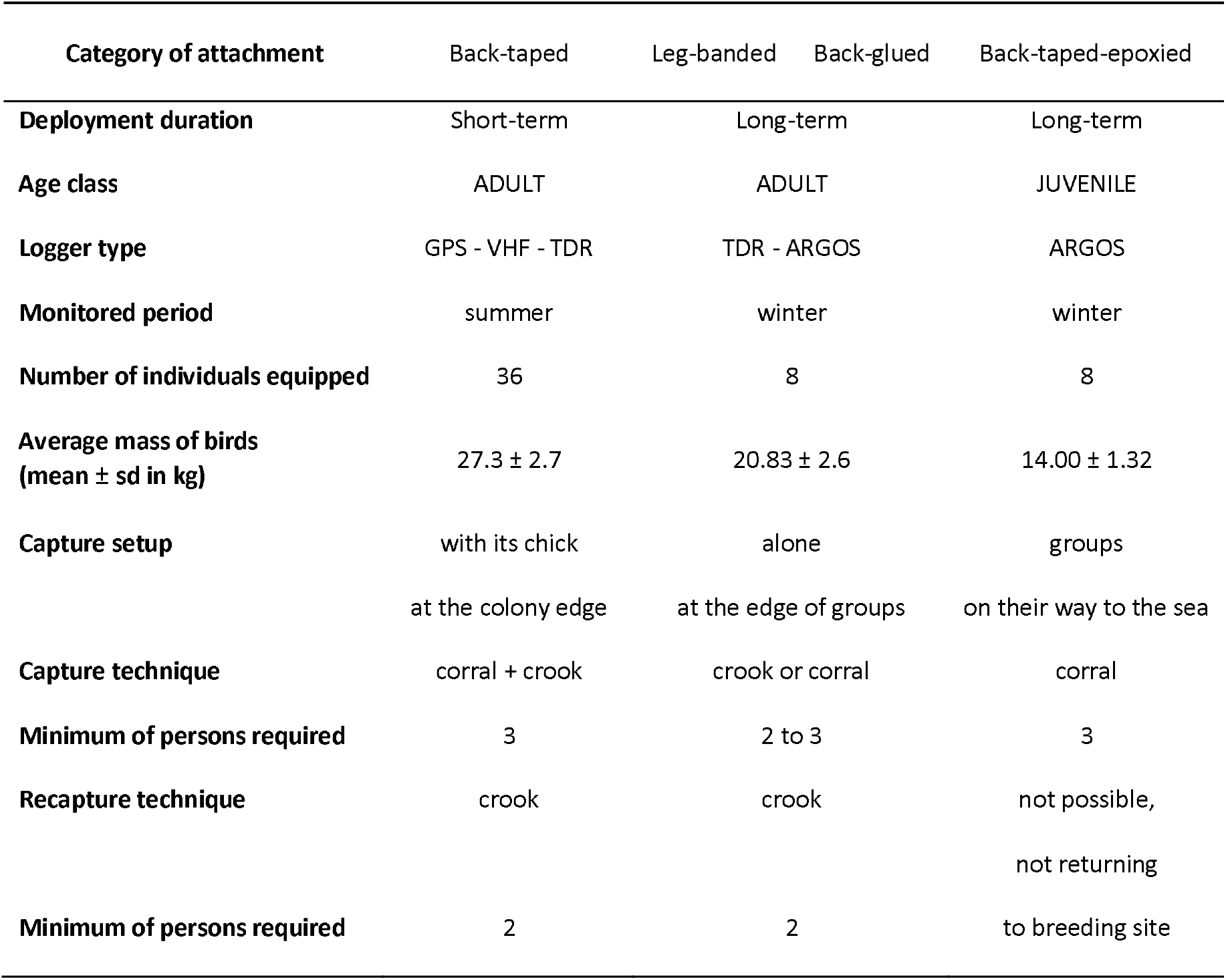
General information on deployments and captures.

| Category of attachment | Back-taped | Leg-banded | Back-glued | Back-taped-epoxied |
| --- | --- | --- | --- | --- |
| **Deployment duration** | Short-term | Long-term |  | Long-term |
| **Age class** | ADULT | ADULT |  | JUVENILE |
| **Logger type** | GPS - VHF - TDR | TDR - ARGOS |  | ARGOS |
| **Monitored period** | summer | winter |  | winter |
| **Number of individuals equipped** | 36 | 8 |  | 8 |
| **Average mass of birds**<br>**(mean ± sd in kg)** | 27.3 ± 2.7 | 20.83 ± 2.6 | | 14.00 ± 1.32 |
| **Capture setup** | with its chick<br>at the colony edge | alone<br>at the edge of groups |  | groups<br>on their way to the sea |
| **Capture technique** | corral + crook | crook or corral |  | corral |
| **Minimum of persons required** | 3 | 2 to 3 |  | 3 |
| **Recapture technique** | crook | crook |  | not possible,<br>not returning |
| **Minimum of persons required** | 2 | 2 |  | to breeding site |

#### a) Short-term deployment: back-taped loggers

We equipped 16 adults in 2017/2018 and 20 adults in 2018/2019 with a GPS-Acc-VHF logger (a combination of a Global Positioning System (GPS), a 3-axis Accelerometer (Acc) and a Very High Frequency (VHF) locator beacon) and a separate Time-Depth Recorder (TDR) (see Additional file 1 for technical details on the loggers). Both are archival devices and, therefore, need to be retrieved to download the data. The VHF locator beacon sends a device-specific signal that allows to locate the equipped birds in the colony and facilitate device recovery. To minimise deleterious effects such as extra drag on diving animals ^20,76^, we followed the recommendations of previous studies. The hydrodynamically-shaped devices represented less than 1% of the penguin’s cross-sectional area, weighed less than 3% of the bird’s mass (Table 1) ^27^ and were attached on the lower back of the birds with adhesive tape.

#### b) Long-term deployment: back-glued loggers

In January 2018 we equipped 8 adult emperor penguins that had completed their moult with an Advanced Research and Global Observation Satellite (ARGOS) platform and a separate accelerometer. ARGOS platforms sent the birds’ location via the Collecte Localisation Satellites (CLS) ARGOS service (Toulouse, France). The streamlined devices were attached by direct contact between cyanoacrylate glue and the feathers in the middle of the lower back of birds ^29,77^.

#### c) Long-term deployment: back-taped-epoxied loggers

In January 2019, we equipped 8 juveniles with ARGOS platforms. The devices were attached to the lower back of birds with adhesive tape that was secured with epoxy glue. Importantly, the epoxy glue did not come in contact with the back feathers.

#### d) Long-term deployment: leg-banded loggers

In January 2018, the same 8 birds equipped with ARGOS loggers (see section b above) were also equipped with an additional TDR sensor that was attached with a leg-bracelet. Similar leg-bracelets had been successfully deployed on other penguin species ^11,30,54,55,78^.

### Capture protocols

A very limited number of scientists have ever handled a non-anaesthetized adult emperor penguin. Handling such an animal can be difficult as they are strong but fragile birds (especially the flippers) with a body mass ranging from 15 (this study) to *ca*. 40 kg depending on age, sex, season and location ^67,79^. While it is always better to transfer such skills directly in the field, this may not always be possible due to the limited number of qualified and experienced persons able to train others. Therefore, our study and the associated protocols aim to fill part of this gap. The techniques developed in this study to approach, capture and handle an adult emperor penguin require, as a minimum, two qualified field staff (referred to hereafter as specialists).

#### 1) Adult-Chick capture protocol

Here, we present a technique to capture an adult emperor penguin with its thermally independent chick during the late chick-rearing period. To avoid larger disturbances, it is ideal to capture birds at the outer rim. Therefore, the first step is to observe the outermost 3-4 rows of animals from a distance, to locate adults that are feeding the same chick several times and that are either stationary or moving towards the outer edge of the colony. Note that allofeeding behaviour is quite common in emperor penguins ^80^ but allofeeders usually do not stay with the same chick at this time of the season (Houstin and Le Bohec, unpublished observations).

#### Capture equipment

Three main tools are required:

– One 2 to 3 m long stick (*e*.*g*. lightweight bamboo sticks) used to direct targeted birds out of the colony (Fig. 1a).

**Fig. 1.**
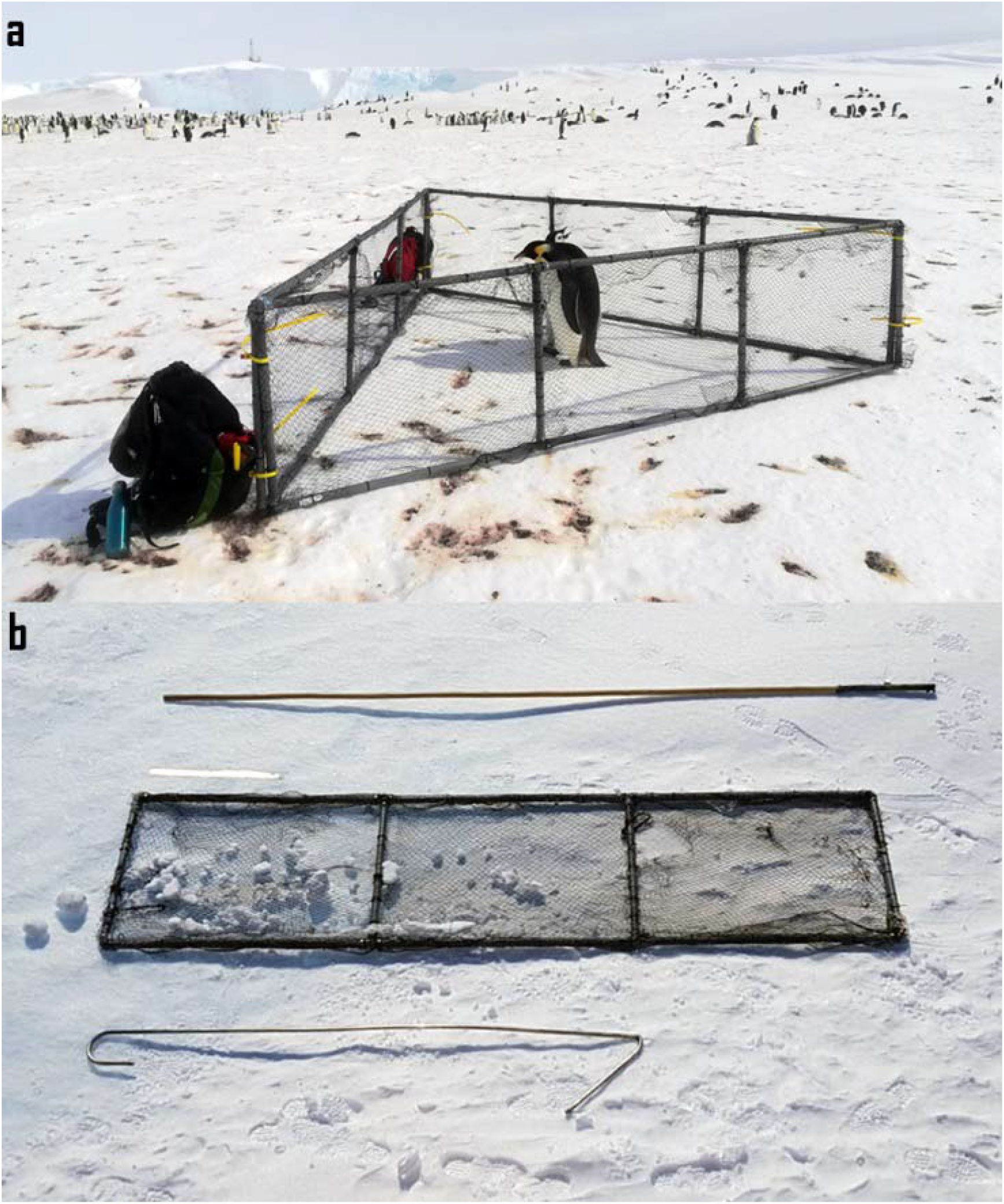
Required tools to capture emperor penguins. **a** An adult-chick pair inside the corral. **b** A 3 m long bamboo stick at the top, one of the panel of the corral (a 50 cm ruler is placed just above it to facilitate scaling) in the middle, and the crook at the bottom. Photo © CNRS-IPHC / CSM.
– One 2.5 m long light crook made of stainless steel or aluminium, bent at 50 cm from one end by an angle of approximately 135° (Fig. 1a), used to direct birds and catch them if necessary. Note that a crook is more efficient than a hook from which penguins manage to escape by twisting their neck.
– A corral made of three separate panels (Fig. 1a). Each panel consists of plastic pipes joined together to form a 3 m by 0.8 m frame. For every meter in length, a vertical plastic tube is added for stability. A polypropylene net (aviary net with a mesh size of 2 cm) is connected to the frame using cable ties. This construction results in lightweight (*e*.*g*. 4.5 kg), sturdy and field serviceable panels. When the panels are connected (Fig. 1b), the triangle formed can be closed with two large reusable cable ties at each of the three joints. We suggest covering one of the panels with a plain fabric, even if this makes the panel more difficult to handle when there is wind. The fabric reinforces the corral, provides shade to the birds and prevents them from attempting to go through the net. It also protects the fieldworkers from wind and allows them to hide behind the panel to calm the birds before release.

#### Corral capture procedure

When the target adult-chick pair is located, the two specialists (one with the bamboo stick, and the other one with the crook) move towards the birds from two sides, starting approximately 40 m away from the colony. The first step for the specialists is to position themselves “behind” the pair, so that the birds are between them and the outer edge of the colony. The second step is to guide the pair slowly out of the colony by walking one-step at a time behind them. The specialists move very slowly to minimise the disturbance of the colony. The resulting disturbance is minimal (Additional file 3) especially if compared to a natural event like the intrusion of a Weddell seal (*Leptonychotes weddellii*) into the colony (Additional file 4). Meanwhile, the two assistants are positioned at a distance of approximately 100 m from the colony edge with the three corral panels and await instructions by radio communication at a minimal volume. Situational awareness is crucial to anticipate the location where the pair will exit the colony and to ensure a fast least-disturbing capture.

Once the pair is ∼30-40 m outside the colony, one assistant hands one of the panels to the specialist with the bamboo stick, and returns to his/her own position. Once the panels have been placed equidistantly (∼30 m) around the penguin pair, everybody moves closer to the pair and close the corral around it. It is to be noted that the last few meters (< 5 m) before the corral is fully closed, the team has to move in a smooth, swift and highly coordinated manner, so that no escape route is presented. If correctly executed, the penguins will remain stationary, looking for the best escape route, and find themselves in the closed corral before an escape is attempted. The specialist with the crook helps to close and secure the corral with reusable cable ties. If the adult attempts to escape, use the crook to catch the bird and prevent the escape (see section 2 -single adult capture protocol). Four persons are the optimal number to carry out this capture protocol. If everybody is experienced, it can be executed comfortably, for the animals as well as for the scientists, with three people. The whole procedure is presented in detail in additional file 3. After capture and manipulation, we recommend to let parent and chick rest and calm down for a few minutes in the corral to increase the chance that they stay together upon release. To release the birds in a particular direction, the cable ties of the edge facing the desired exit direction are unzipped and the corral sufficiently opened to allow the birds to go through (Additional file 5).

We used this method to capture a pair of adult with its not-yet-moulted chick to increase our chances of recapture. Indeed, breeders with moulted chicks ready to fledge or with chicks having sufficient reserves to perform their moult and fledge on their own are more prone to end their breeding cycle, defined by the “abandon” of their chick ^56^. Devices were recovered after one to three foraging trips.

#### 2) Single adult capture protocol

Two techniques can be used to capture a single adult emperor penguin; the choice depends on the behaviour of the bird while approaching, the availability of assistants, and the weather conditions. As described above for the pair of an adult and its chick, the corral can be used to trap a single adult in a very similar way. Nonetheless, due to the fact that solitary birds are more mobile and usually more vigilant to their environment the corral method may be difficult, which is especially true during heavy winds or a blizzard.

An alternative and efficient technique is to use a crook to catch the bird (Additional file 6) as explained in Cockrem et al. ^58^. The crook capture requires two people and in contrast to the corral protocol and the deployment of loggers can also be conducted in bad weather. Once the bird is isolated, one specialist places the crook around the neck of the bird preventing the penguin to escape by tobogganing, *i*.*e*. moving on its belly. Meanwhile, the other specialist grabs the tibiotarsi of the bird and holds them firm. The crook is gently removed and placed away from the capture site, and the penguin secured by the two specialists, one in front of the bird and one at the back. The crook-carrying specialist should be carrying the necessary supplies for manipulation in his/her backpack, because, after the capture, he/she will have his/her hands free, while the other specialist is still holding the bird.

We used this technique to recapture adults for device recovery or to capture non-breeding (*e*.*g*. moulting or post-moulting) adults.

#### 3) Fledging juvenile capture protocol

For their first departure at sea, juvenile emperor penguins usually leave the colony in small groups. A group capture with the corral is, therefore, more efficient and potentially less stressful for the birds. The protocol is similar to the adult-chick-pair capture, but here an entire group of juveniles is slowly encircled by three corral bearers. As emperor penguins are social animals, it is likely that keeping the group together reduces the stress of manipulated individuals and facilitates the remainder of their travel towards the sea after release. Juveniles of interest are removed individually from the corral for the manipulation and returned afterwards. All juveniles are released together after all target animals have been handled.

### Adult emperor penguin handling protocol

Similarly to Cockrem et al. ^58^, and as shown in the Additional file 7, the bird is caught and maintain upright by one specialist (S1). Once the bird is secured in this position, the second specialist (S2) approaches and bends the penguin’s head towards the ground while S1 grabs the legs above the ankles to lay the bird on its belly. When the bird is lying on the ground, S2 kneels over the bird with its head between (below) the legs of S2. In this position, the bird is immobilised. It is crucial that the flippers, the most fragile part of the bird, are unrestrained and untouched, throughout the whole process. If an assistant is available, he/she can hold the legs of the birds and stretch them (foot soles pointing towards the sky). Working with three people allows S1 to deploy the loggers seated next to the penguin and reduces manipulation time. Most penguins stay quiet in this position with some few second long bursts of intense activity: a gentle but firm pressure on the back and pulling the foot soles upper and further from the ground helps to calm the bird.

### Equipment protocols

During manipulation, the bird’s eyes were always covered with a hood to reduce stress level ^29^ and birds were handled at distance from the edge of the colony to avoid conspecifics’ disturbance (usually > 40 m, thus well above the 5 m limit recommended in the General guidelines produced by the Antarctic Treaty Consultative Meeting ^81^).

#### a) Short-term deployment: back-taped loggers

Before starting with the attachment, we used a cardboard stencil and waterproof tape that is a bit larger than the logger to demarcate the precise location of the equipment and the placement of the strips of adhesive tape on the penguin (see this in detail in Additional file 8). Following studies from Wilson et al. ^77,82^ and numerous subsequent short-term studies on other penguin species ^83–87^, we used a rounded knife to lift a few feathers from the back of the penguin and insert pre-cut strips of waterproof adhesive tape (*e*.*g*. Tesa® tape 4651, Beiersdorf AG, Hamburg, Germany). To further reinforce the attachment, we added glue (*e*.*g*. cyanoacrylate glue, Loctite 401, Loctite, Henkel AG., Düsseldorf, Germany) between the adhesive part of the tape strips and the logger. Cable ties (*e*.*g*. Panduit, Panduit Corp, Illinois, USA) should be tightened with a cable tie gun. For a deployment period of more than one month, we recommend to add glue on top of the tapes. After manipulation was completed, we marked the bird before release with a hair-dye painted number that will last until the following moult (*e*.*g*. Schwarzkopf, Palette dark-blue N°909, Henkel AG., Düsseldorf, Germany, see Additional file 9 to behold a marked bird).

We used this technique to deploy GPS-Acc-VHF (Axy-Trek from TechnoSmArt) and TDR (g5+ from Cefas) devices (see Additional file 1 for technical details and Fig. 2a to view an equipped bird) on adult emperor penguins at the end of their breeding cycle (see respective movies of deployments in Additional files 8 and 10).

**Fig. 2.**
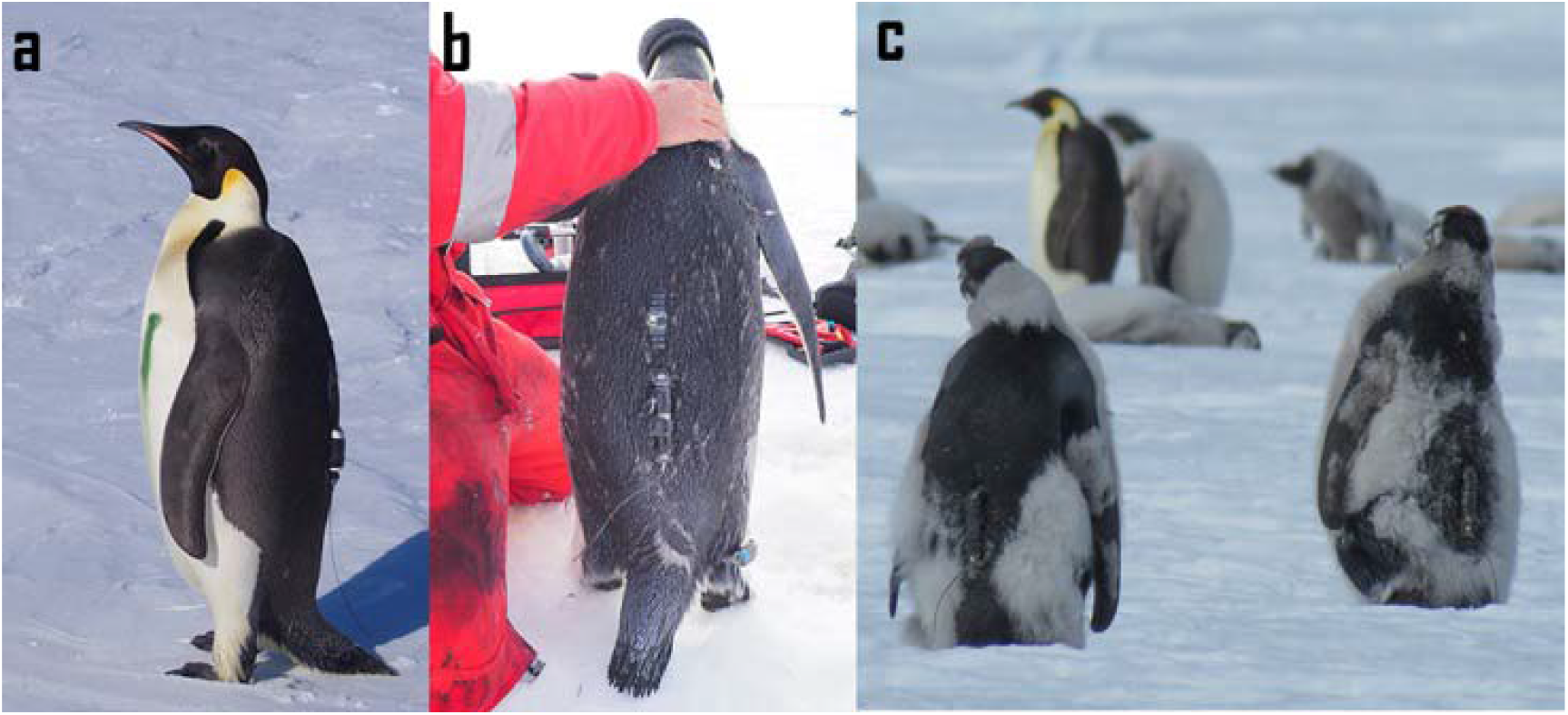
Pictures of the different deployments performed. **a** Adult emperor penguin equipped with back-taped loggers (a TDR in the middle of the back and a GPS underneath). The green line on the bird’s belly is non-permanent marking. **b** Adult emperor penguin equipped with back-glued loggers (an accelerometer in the middle of the back and ARGOS satellite transmitter underneath) and a leg-banded logger on its right foot. **c** Juvenile emperor penguins both equipped with a back-taped-epoxied logger. Photo © CNRS-IPHC / CSM.

#### b) Long-term deployment: back-glued loggers

According to protocols from previous studies ^40–42^, that conducted long-term deployment of biologgers on the back of penguins, we fixated the lower side of the loggers directly to the feathers using cyanoacrylate glue (Loctite 401). The attachment was secured with polyamide cable ties as described above (Fig. 2b).

We used this technique to deploy Spot-367 ARGOS loggers from Wildlife Computers and WACU accelerometer from MIBE-IPHC-CNRS on adults close to finishing their moult (see Additional file 11 to identify an emperor penguin in that stage).

#### c) Long-term deployment: back-taped-epoxied loggers

Similar to the short-term deployment protocol, the logger is attached to the feathers using pre-cut lengths of Tesa® tape on the entire logger length (sparing exposed sensors if any). The overlap between tape strips is reinforced with cyanoacrylate glue (Loctite 401). We used two polyamide cable ties around the head and one at the bottom of the logger to secure the attachment. The supernumerary cable tie on the head is added for extra safety. Once all the adhesive strips and cable ties were fixated, we applied epoxy adhesive (Loctite EA 3430) on the mounting (sparing exposed sensors if any) to reinforce the waterproofness and robustness, adapting methods from other studies ^88,89^. The attachment procedure is shown in the Additional file 12.

We performed this deployment on fledging chicks. We selected the individuals most advanced in their moult, *i*.*e*. presenting no more down on their back (Fig. 2c). The lower survival rate of the juveniles during the first year at sea compared to adults ^90^, their non-return to breeding colonies before several years ^72^ and their unfinished growth, prevent the use of other types of externally attached devices.

#### d) Long-term deployment: leg-banded loggers

To reduce drag and behaviour disturbance induced by devices on the back of penguins, we developed a leg-band (bracelet) for mounting TDR-loggers on emperor penguins. We designed two similar types of bracelet, a first version that we deployed (Fig. 2b), and a second version incorporating slight changes and improvements. A detailed manual of the mounts is provided in Additional file 13. We designed the bracelet to mount a Lotek Lat 1800 TDR (see Additional file 1 for specifications) but the bracelet can be easily adapted to other types of TDR.

The TDR is fixed to a rubber cable tie (Panduit, ERTM-C20) covered with heat-shrinkable sheath and attached around the bird tibiotarsus by closing the cable tie just above the ankle, like a bracelet. A built-in lock prevents the cable tie to tighten itself after deployment. The bracelet fits loosely with ∼1 cm space between the bracelet and the leg. The shape of emperor penguin’s legs prevents the bracelet from spinning around the leg. When properly set up and attached (Additional file 13), the attachment does not interfere with egg or chick placement on the bird feet during the breeding season. Deployment time lasts less than 3 minutes. On retrieval, the bracelet is easy and quick to remove (within a few seconds) by cutting the rubber cable tie with pliers.

## Results

### a) Short-term deployment: back-taped loggers

In 2017-2018, 16 deployments were performed: 10 between November 27^th^ and December 2^nd^, of which 6 devices were recovered and redeployed between December 10^th^ and 12^th^. None of the devices of the second deployment session were recovered, resulting in 38% recovery. In 2018-2019, 20 deployments were performed, 10 between November 05^th^ and 07^th^, which all were recovered and redeployed between November 25^th^ and December 6^th^. Six devices of the second deployment were recovered, resulting in 83% recovery. We conducted intense VHF and visual (binocular) surveys for equipped birds (approx. every 4 hours), thus we are confident that we retrieved all the loggers from returning birds. All VHF units of recaptured birds were working and unequipped birds have been regularly identified afterwards by their hair-dye painted number on their chest.

Bird feathering on recovery was intact and no physical damage on the bird or on the device was apparent. All loggers were still securely attached, even after the longest deployment of 25 days (Table 2). Our recovery rate for November (90%) is similar (z-test, p-value > 0.05) to those of previous studies (Table 2). The recovery rate from December 2018 (30%), despite being higher, is statistically similar to what Robertson ^56^ recorded for deployments performed in December on the opposite side of Antarctica (near Australia’s Mawson Station) with a loss rate of 89%. The probability to recover a device deployed in December is significantly lower (z-test, p-value < 0.05) than in November.

**Table 2.**
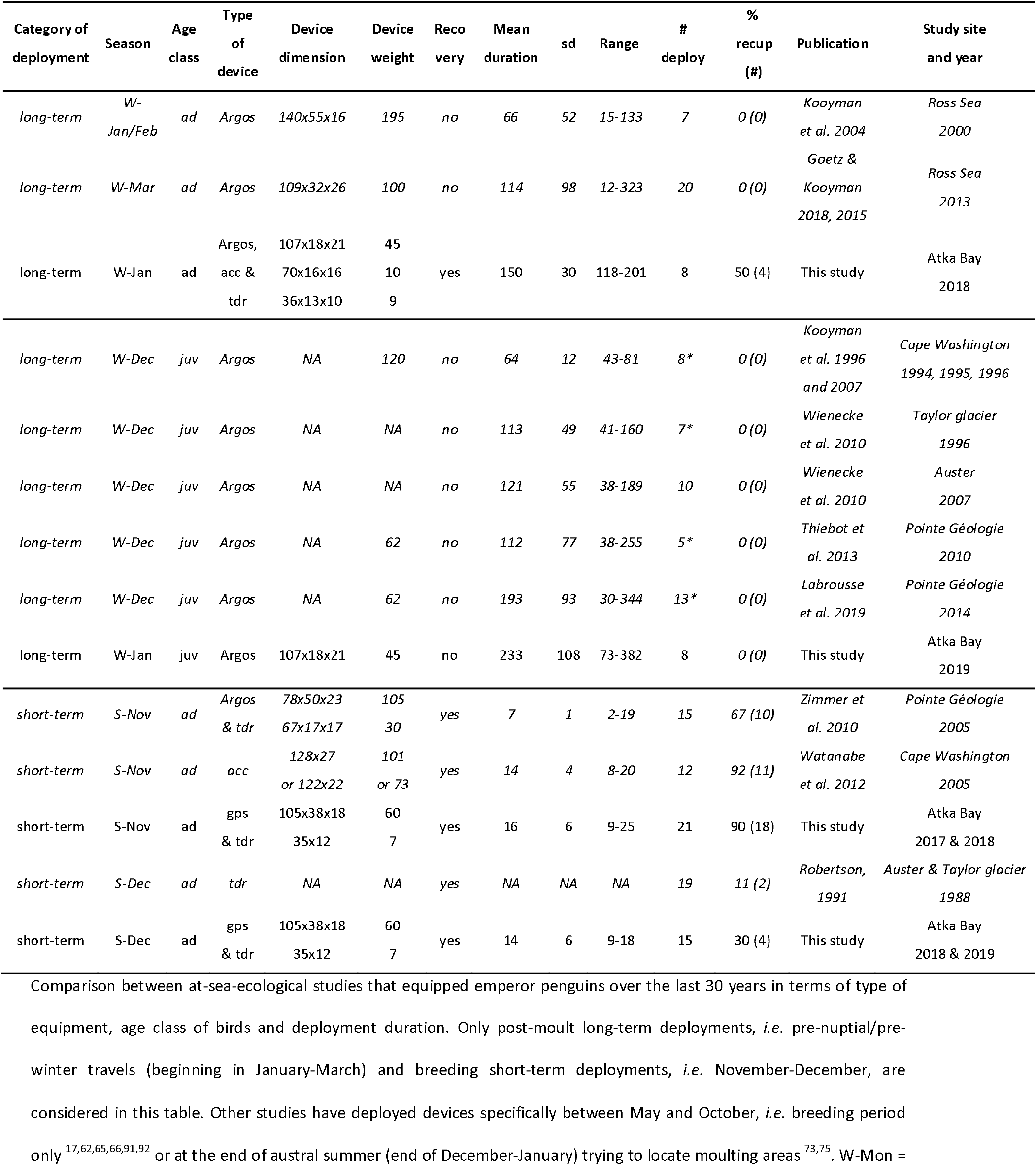
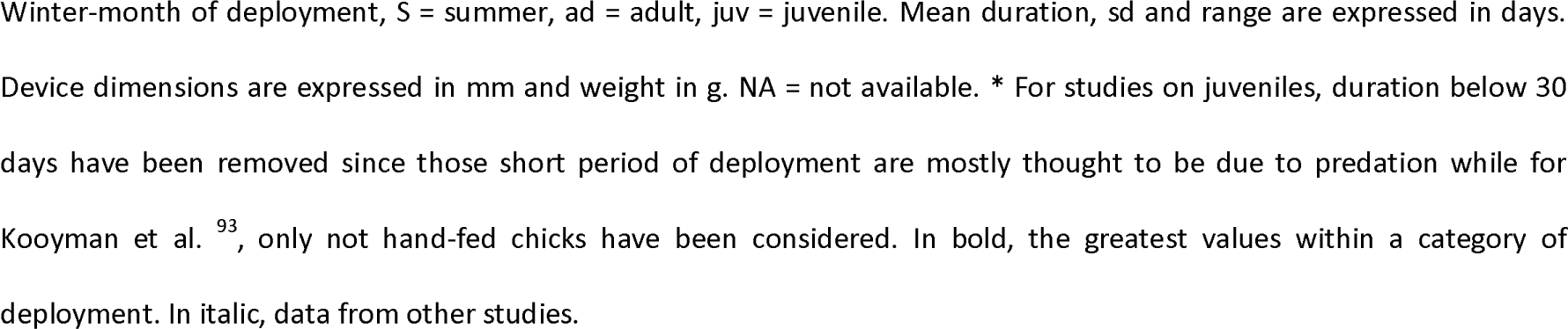
Comparison between at-sea-ecological studies that equipped emperor penguins over the last 30 years.

### b) Long-term deployment: back-glued loggers

Our study is the first to report recapture of emperor penguins after a whole-winter deployment (January to November). Identified by the number painted on their chest (Additional file 9), 4 of the 8 birds equipped in January 2018 were resighted and recaptured in November 2018 (Table 2). All of them had lost the devices on their back. Instead, there was a line of missing/broken feathers (Fig. 3). No injury was detected. Signals from all ARGOS devices were lost during the winter. The mean transmission period was 150 ± 30 days (range 118-201 days, Table 2), significantly exceeding the previous average deployment durations of 66 ^42^ (p-value > 0.05, ANOVA) and 114 days ^43^ (p-value < 0.05, ANOVA) from all previous similar studies.

**Fig. 3.**
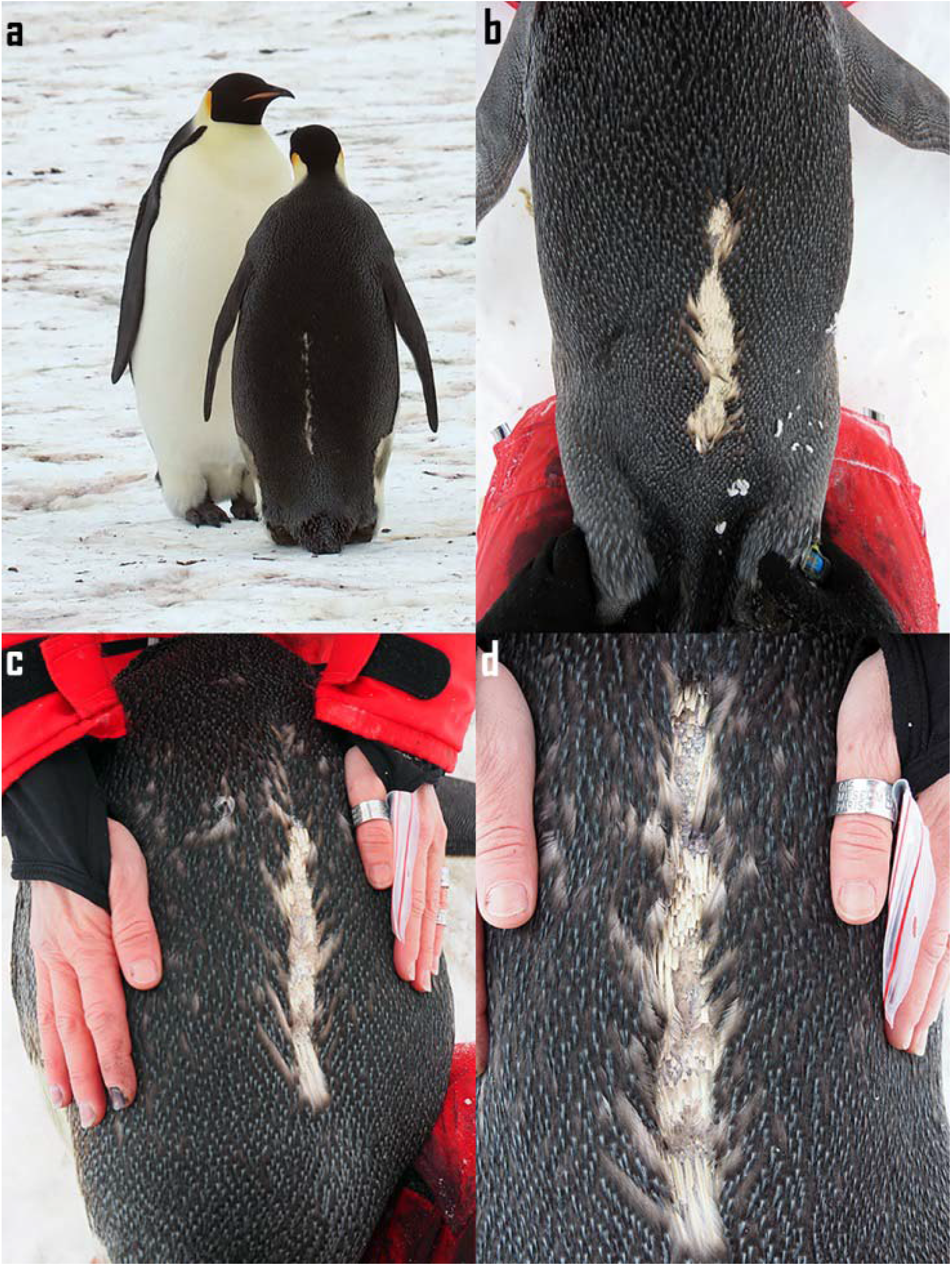
Back of the 4 penguins having lost their back-glued loggers during the winter. Photo © CNRS-IPHC / CSM.

### c) Long-term deployment: back-taped-epoxied loggers

Three of the 8 juveniles equipped in January 2019 transmitted until their annual moult in January 2020. None of the birds did return to their native colony for moult; an observation congruent with the conjecture that juveniles of 1.5-years of age do not come back to moult at their birth colony for the first year ^44,72^. This tracking period of a full year, from January 2019 to January 2020 (Table 2) is the longest documented deployment duration for the genus Aptenodytes. None of the 5 remaining birds were spotted on colony despite visual search in summer 2019/2020. One device stopped transmitting after 73 days while the four others lasted between 142 and 185 days resulting in an average deployment duration of 233 ± 108 days. This mean deployment duration is longer than any previously reported (p-value > 0.05, ANOVA, see Table 2 for mean ± sd values) but not significantly compared to Labrousse et al. ^44^ (p-value < 0.05, ANOVA).

### d) Long-term deployment: leg-banded loggers

Our study is the first to perform a year-round deployment and data collection on emperor penguins. The 4 adults recaptured without their back-mounted loggers were still carrying their leg bracelet mounted TDR, providing an unprecedented record of an entire year of high frequency (1 Hz) depth and temperature logs for emperor penguins.

For all recaptured birds, the leg-bracelet mounting did not present any damage, the bracelet and the TDR were at the same position of their deployment, suggesting that the device did not rotate around the leg during the deployment period. However, all recaptured birds had lost a few feathers especially on the inside part of the leg and showed signs of abrasion in the form of a slight reddening of the skin and peeling under the bracelet area. Two of them had small sore spots on their tarsi. No limping was observed before or after removal. An illustrated comparison between an equipped and an unequipped leg after recovery can be found in the Additional file 14.

## Discussion

To our knowledge, two of the four deployment methods presented in this study are new developments for this species. These two methods allowed for the longest documented deployment duration for this species. The description of those methods, paired with an exhaustive documentation, aims to facilitate and enhance future research on this species.

### Capture and handling

All capture techniques presented in this study yield minimal colony disturbance regardless of the period of the breeding cycle. The described handling is safe for birds and handlers, and only a minimal number of trained personnel is required. We recommend the use of the corral if no member in the field team is accustomed to handling a crook or a hook on at least one penguin species.

### Deployments

#### a) Short-term deployment: back-taped loggers

Our study is the first to report the deployment of GPS devices on emperor penguins. GPS devices have already been deployed on penguins ^87,94^, yet never on emperor penguins on which only ARGOS devices ^62–64,95^ or dive-pattern-analysis related loggers ^56,64,95^ have been deployed during the late chick-rearing period, presumably due to a combination of technical and logistical constraints. The methods presented herein allow the deployment of high-resolution data acquisition loggers with a high probability of recovery once the phenology of the colony has been assessed, for instance by the size and moulting stage of chicks. At Atka Bay, the best deployment period is in November with a logger recovery rate of 90%. The low recovery rate (38%) during the 2017-2018 season can be explained by logistical issues we encountered and not the deployment technique. An unexpected late on-site arrival led to late deployment of 11 loggers in December 2017, compared to 4 in December 2018, and consequently to a substantial loss of devices.

To optimise the recovery rate of devices deployed at the end of the breeding season, we recommend to deploy devices on adults with medium-sized chicks at the very beginning of chick moult. Supported by the secure attachment of the presented technique, we furthermore suggest increasing deployment time rather than to recover loggers and redeploy them.

#### b) Long-term deployment: back-glued loggers

Our study is the first to document the recapture of a long-term equipped emperor penguin and thus able to assess (i) the state of the bird, (ii) the state of devices, and (iii) provide evidence explaining the loss of signal from communicating-satellite-relayed loggers reported in previous studies. Until now, five studies had performed long-term deployments on emperor penguins right after the moult (Table 2), all using ARGOS platforms and cyanoacrylate glue to attach the logger directly to the back-feathers of the birds. None of the birds were resighted, perhaps partly due to the logistical possibilities to reach the colony in the following years at other study sites. Two sets of deployments were made on adults ^42,43^, three on chicks ^40,41,44^.

Our results show that both glued devices, the ARGOS transmitter and the small accelerometer, which vary in size and weight (Additional file 1), were lost in the same manner on all birds. We speculate that the cyanoacrylate glue rigidifies the feathers, which then become brittle and break with either the continuous birds’ movements and/or their attempts to remove the device. Wilson et al. ^77^ also observed this device-sized hole in the feather layer after winter deployment on four Magellanic penguins (*Spheniscus magellanicus*) using epoxy instead of cyanoacrylate glue. In our study, five of the eight ARGOS signals were lost while birds had been on fast ice at the breeding site for several weeks. The longest duration of deployment (201 days, Table 2) was recorded for the only bird that did not spend any extended time on ice. When the birds remain on the fast ice at their breeding colony, they are exposed to extremely cold temperatures, below -50°C in Atka Bay ^96^. We assume that such cold temperature could have two different effects. Either the cyanoacrylate induced brittle feathers become very weak when exposed to such cold temperatures and, therefore, are easier to break during the penguins’ movement, or the brittle feathers do not provide the proper isolation against the cold, thus altering the heat transfer and/or thermoregulation of the bird (as observed on pup grey seals (*Halichoerus grypus*) by McCafferty et al. ^97^). This could lead to the animal preening their feathers fiercely and thereby removing the logger by breaking/removing the brittle glued feathers. In addition, blizzards may have caused the accumulation of ice around the loggers, amplifying the above effects and tearing the device off ^42,45^. However, we suggest that the huddling behaviour of emperor penguins ^67,98^ would prevent ice accumulation in the middle of the winter. Another possible explanation could be the timing of deployment. Devices were attached just at the end of the moult, a time when feathers may not yet be fully developed despite a meticulous bird selection process (Additional file 10). Their growth after deployment could potentially have added some slack and thus reinforced the pull on the feather shafts, ultimately leading to their breakage after few months. The loss of back feathers undoubtedly leads to a diminution of insulation that causes a greater heat loss. The resulting increase in energetic needs reduces fasting capabilities and forces the birds to compensate by finding more prey items when they return at sea to forage in order to replenish their reserves while accumulating food for their chick. As body reserves management is critical, especially for this species, any significant heat loss is likely to impact breeding success.

In tagging procedures, the ethical principle of *Refinement* from the Three Rs ^32^, *i*.*e*. the use of methods which decrease any adverse effect, should apply. It is likely that the device loss we recorded also happened during previous deployments that used this technique. We consider that the loss of the device and resulting consequences for the birds are unacceptable and that our findings combined with the ones from Wilson et al. ^77^ should prevent further use of glue directly on the birds feathers for long-term deployment on penguins as it is currently practiced. We propose an alternative technique that has not yet been shown to induce such damage. The tape does not include acrylamide glue and is therefore less likely to brittle the feathers.

#### c) Long-term deployment: back-taped-epoxied loggers

To investigate the distribution at-sea according to the age-class we employed the back-taped-epoxied technique on juveniles. So far, this age-class had only been tagged using the back-glued method (Table 2).

Three of the juveniles (40%) retained their device for an entire year, thus achieving the longest duration of back-mounted logger deployment possible in penguins. The previous longest durations recorded for juvenile emperor penguins were of 344, 298 and 271 days ^44^ with one bird (6% of the deployments) approaching the one year length duration. The mean duration of our long-term deployments (233 ± 108 days) with the taped-epoxied technique is longer than any previously reported on juvenile or adult emperor penguins (Table 2). Therefore, we are confident that the technique presented is a significant improvement for tracking of penguins and understanding their activities at sea, even if the contribution of a possible gain resulting from the evolution of technologies is not measurable.

Some of the previous studies using glue on juveniles approached long-term attachment duration, with only one of the 48 juveniles exceeding 10 months of equipment. The lack of recovery of ARGOS devices deployed on juveniles can be explained (i) by the fact that the birds moult outside their original colony ^62,73–75^, or by the loss of devices as suggested by our results on adults. Electronic failure or bird predations are also alternative hypotheses ^42^. Nevertheless, in addition to the possible loss of feathers and insulation previously discussed, the glue has the potential to cause thermal skin burns ^99^. Juveniles are more vulnerable than adults as their foraging skills (including their ability to dive, to capture prey, and to find productive feeding grounds) are not yet fully developed, and their experience to escape predators is also minimal ^100–102^. The additional cost mentioned above induced by a glued device may negatively impact the survival of the juveniles during their first months in their new marine environment that they experience for the first time. As a result, for studies requesting the deployment of back-mounted devices on penguins for long-term duration, the use of glue on feathers should be entirely avoided, and we recommend to use instead a mix of Tesa® tape strips (feathers’ side) and epoxy (on the strips covering the device) to reinforce adhesion.

We could not show that this attachment will last on adult emperor penguins as long as for juveniles. An early departure from the field due to logistical constraints prevented us to deploy this new technique on fully moulted adult emperor penguin. Adult emperor penguins experience very harsh environmental conditions on the sea ice, especially at their breeding site, during winter with temperatures below -50°C and wind speeds above 150 km/h at Atka Bay ^96^. Average deployment durations on adults are less than six months (Table 2) and need to be improved to cover the entire breeding cycle and justify the impacts on the birds’ welfare. We are convinced that new techniques should be tested such as the promising one presented in this study for juveniles.

#### d) Long-term deployment: leg-banded loggers

We developed and tested a leg-band TDR mount to enable year-round deployments on adult emperor penguins. The deployed leg-band TDR mount collected high frequency (1 Hz), pressure and temperature data during the whole year. This data will allow a detailed analysis of foraging activities and water column exploitation over a full year for the same birds.

Leg-band mounted devices had already been deployed on penguins ^11,30,54,55,103–105^, but not yet on the emperor penguin species, which can walk over long distances on sea ice ^65^. Often, the condition of the birds at retrieval are not mentioned, however, some of the studies reported similar leg irritations ^30,105^ (Raclot personal communications; Houstin, Fournier and Le Bohec, unpublished observations) as the ones we observed in this study. Such irritations might be due to the fact that emperor penguins can walk over long distances on sea ice to reach the water ^65^. The commonly accepted flying bird banding technique is also known to cause unintentional damage like sores, inflammation, or even loss of feet in extreme cases ^106–108^, thus the irritations observed here can be considered as a minor impact.

We suggest that the use of non-continuous heat-shrink tubing and the glue around the head of the rubber cable tie created a small ledge in the otherwise smooth surface that irritated the birds’ leg-skin. From this observation, we have designed an improved version for the second season (Additional file 13), which could not be tested due to early departure from the field site. The continuous heat-shrinkable sheath in the updated bracelet attachment will likely reduce friction between the leg and the bracelet and ideally avoid skin irritation. If feathers are still lost, we expect the tibiotarsus to be less irritated and the occasional development of sores prevented. To prevent the formation of glue flakes, glue will only be applied inside the cable tie’s closure, with parsimony, and not around the whole head.

At retrieval, the mounting did not show any damage or sign of wear and is expected to last several years before the elastomeric cable tie breaks. Consequently, before deploying such a system, a strategic plan for its retrieval is crucial. Emperor penguins are non-nesting seabirds, breeding freely on sea ice within a mobile colony ^109^, makes the recapture of birds difficult, especially after more than one year when the annual moult removed any externally painted-markings. Due to their PIT-tag, all birds manipulated can be life-long identified by automatic Radio-frequency identification (RFID) detection systems, without requiring recapture or visual observation ^110–112^ and without long-term deleterious effects of flipper-banding for life ^35,37,38^. Such automatic detection systems have been successful at detecting emperor, Adélie and king (*Aptenodytes patagonicus*) penguins over the last years at the Pointe Géologie archipelago and Crozet and Kerguelen archipelagos (Le Bohec, Houstin, Chatelain and Courtecuisse, unpublished observations, ^113^). Such a system will be deployed at Atka Bay colony starting in summer 2021-2022 to improve our retrieval ratio of 50%, compared to the retrieval rate for nesting birds that is between 60 and 90% ^11,30,104,105,114,115^. This technological improvement will allow in the years to come to recapture birds even after several years of deployment like for nesting birds ^30^ (Raclot personal communications; Houstin and Le Bohec, unpublished observations).

Specifically, with this bracelet technique, multi-year deployments might be considered. Scientific programs running in Antarctica are not always able to return several years in a row, and this technique of deployment offers some flexibility. Solutions still need to be developed for communicating devices that need to get out of water regularly to send or receive telemetry (GPS, ARGOS) or for biologgers that would record too noisy data when positioned on the leg. However, for small data loggers, that are able to record environmental variables (*e*.*g*. hydrostatic pressure, water conductivity and temperature, environmental luminosity) on a multi-year scale ^102,116^, the leg-band technique appears as a high potential alternative.

## Conclusion

Ethical concerns raised by the use of measuring devices on wild animal are not new ^28^ and a recent review ^117^ addressed the current pros and cons on attachment issues. To ensure data is of exemplary quality from a scientific and ethical point of view, the potential deleterious effects of deployment procedures (capture-attachment-recapture) must be assessed and mitigated. Our study provides highly detailed procedures to capture/recapture and externally attach telemetry devices on emperor penguins. We, therefore, consider this study as a significant advancement by (i) stating clearly the impact of using glue for biologging device attachment on emperor penguins, (ii) helping to assess long-term loggers loss reasons (notably ARGOS transmitters), (iii) presenting two promising attachment techniques of biologging devices on emperor penguins in detail, and (iv) explicitly providing techniques to capture and handle emperor penguins with a limited amount of disturbance as well as a maximum of safety and efficiency. This publication is intended to serve as a resource to facilitate future research on this iconic species.

This study aims also to encourage researchers and journals to give more exposure to fieldwork methodology in scientific publications not specifically methodology oriented, in particular to techniques developed and tested but not successful in the field. We are convinced that too much time and resources are allocated to the development of techniques already tested but not shared because of their failure. Tests, errors and failures are inherent of research and should be, to some extent, valued as significant results; a practice that would benefit to both scientists and animals.

## Abbreviations

GPS: Global Positioning System
TDR: Time-Depth Recorder
ARGOS: Advanced Research and Global Observation Satellite
MPA: Marine Protected Area
PIT: Passive Integrated Transponder
VHF: Very High Frequency
Acc: Accelerometer
CLS: Collecte Localisation Satellites
GLS: Global Location Sensor
RFID: Radio-frequency identification.

## Declarations

### Ethics approval and consent to participate

The AWI long-term program “MARE” (*Monitor the health of the Antarctic maRine ecosystems using the Emperor penguin as a sentinel*), to which this study belongs, and all procedures were approved by the German Environment Agency (Umweltbundesamt-UBA permit no.: II 2.8 – 94033/100 delivered on the 04/10/2017 and 04/10/2018), and conducted in accordance with the Committee for Environmental Protection (CEP) guidelines.

## Consent for publication

Written informed consent for publication was obtained.

## Availability of data and material

All data generated or analysed during this study are included in this published article and its supplementary information files will be available in the “Publishing Network for Geoscientific & Environmental Data” (PANGAEA, https://www.pangaea.de/).

## Authors’ contributions

AH and CLB conceived the ideas and designed the methodology and protocols. AH, CLB, DZ, SR, BF, AW conducted fieldwork. AH, DZ and CLB led writing of manuscript. VPB, DC, AA, JF gave conceptual advice. All authors contributed substantively to the drafts and gave final approval for publication.

## Acknowledgements

We thank the Alfred Wegener Institut Helmholtz-Zentrum für Polar-und Meeresforschung, Logistics Department, the overwinterers and campaigners at Neumayer Station III for their invaluable support in 2017-2018 and 2018-2019 seasons.

We are very grateful to Prof. Patrick Rampal for support in initiating the project, to Dr. Laurent Godet (CNRS) for lending his VHF antennas and receivers, and to Dr. Olaf Eisen (AWI) for his invaluable help with fieldwork and related activities.

We warmly thank ‘Galileo’; and ‘ProSiebenSat.1 TV Deutschland GmbH’ for their shooting in the field and providing Additional files 3 and 5.

## Additional files list

Additional file 1 (.pdf): Table S1. General information on the loggers deployed including the logger type, name, dimensions and weight, as well as the manufacturers name and location

Additional file 2 (.mp4): Movie S1. PIT-tagging of an adult emperor penguin

Additional file 3 (.mp4): Movie S2. Capture of an adult-chick pair

Additional file 4 (.mp4): Movie S3. Weddell seal disturbance on the emperor penguin colony

Additional file 5 (.mp4): Movie S4. Release of an adult-chick pair

Additional file 6 (.mp4): Movie S5. Capture of a single adult with the crook

Additional file 7 (.mp4): Movie S6. Adult emperor penguin handling

Additional file 8 (.mp4): Movie S7. Back-taped GPS deployment

Additional file 9 (.pdf): Fig. S1. Marking of an emperor penguin with a painted number

Additional file 10 (.mp4): Movie S8. Back-taped TDR deployment

Additional file 11 (.pdf): Fig. S2. Adult emperor penguin at the end of its moult

Additional file 12 (.mp4): Movie S9. Back-taped-epoxied logger deployment -epoxy spreading

Additional file 13 (.pdf): Slideshow S1. Leg-banded TDR mounting manual

Additional file 14 (.pdf): Slideshow S2. Leg-banded TDR deployment and recovery

## REFERENCES

1. Rutz, C. & Hays, G. C. New frontiers in biologging science. Biology Letters 5, 289–292 (2009).

2. Marker, L. L., Dickman, A. J., Mills, M. G. L., Jeo, R. M. & Macdonald, D. W. Spatial ecology of cheetahs on north-central Namibian farmlands. Journal of Zoology 274, 226–238 (2008).

3. Kays, R., Crofoot, M. C., Jetz, W. & Wikelski, M. Terrestrial animal tracking as an eye on life and planet. Science 348, aaa2478–aaa2478 (2015).

4. Fortin, D. et al. Wolves influence elk movements: behavior shapes a trophic cascade in yellowstone national park. Ecology 86, 1320–1330 (2005).

5. Ropert-Coudert, Y., Beaulieu, M., Hanuise, N. & Kato, A. Diving into the world of biologging. Endangered Species Research 10, 21–27 (2009).

6. McIntyre, T. Trends in tagging of marine mammals: a review of marine mammal biologging studies. African Journal of Marine Science 36, 409–422 (2014).

7. Bograd, S., Block, B., Costa, D. & Godley, B. Biologging technologies: new tools for conservation. Introduction. Endangered Species Research 10, 1–7 (2010).

8. Clarke, J. et al. Sex differences in Adelie penguin foraging strategies. Polar Biology 20, 248– 258 (1998).

9. Sato, K. et al. Buoyancy and maximal diving depth in penguins: do they control inhaling air volume? The Journal of experimental biology 205, 1189–1197 (2002).

10. Wilson, R. P. et al. Remote-sensing systems and seabirds: their use, abuse and potential for measuring marine environmental variables. Marine Ecology Progress Series 228, 241–261 (2002).

11. Bost, C. A., Thiebot, J. B., Pinaud, D., Cherel, Y. & Trathan, P. N. Where do penguins go during the inter-breeding period? Using geolocation to track the winter dispersion of the macaroni penguin. Biology Letters 5, 473–476 (2009).

12. Weimerskirch, H. et al. Lifetime foraging patterns of the wandering albatross: Life on the move! Journal of Experimental Marine Biology and Ecology 450, 68–78 (2014).

13. Chimienti, M. et al. Taking movement data to new depths: Inferring prey availability and patch profitability from seabird foraging behavior. Ecology and Evolution 7, 10252–10265 (2017).

14. Pistorius, P. et al. At-sea distribution and habitat use in king penguins at sub-Antarctic Marion Island. Ecology and Evolution 7, 3894–3903 (2017).

15. Massom, R. A. et al. Fast ice distribution in Adélie Land, East Antarctica: Interannual variability and implications for emperor penguins Aptenodytes forsteri. Marine Ecology Progress Series 374, 243–257 (2009).

16. Zimmer, I., Wilson, R. P., Beaulieu, M., Ancel, A. & Plötz, J. Seeing the light: Depth and time restrictions in the foraging capacity of emperor penguins at pointe géologie, Antarctica. Aquatic Biology 3, 217–226 (2008).

17. Ancel, A. et al. Foraging behaviour of emperor penguins as a resource detector in winter and summer. Nature 360, 336–339 (1992).

18. Hussey, N. E. et al. Aquatic animal telemetry: A panoramic window into the underwater world. Science 348, 1255642–1255642 (2015).

19. Carter, M. I. D., Bennett, K. A., Embling, C. B., Hosegood, P. J. & Russell, D. J. F. Navigating uncertain waters: a critical review of inferring foraging behaviour from location and dive data in pinnipeds. Movement Ecology 1–20 (2016). doi:10.1186/s40462-016-0090-9

20. Culik, B. & Wilson, R. P. Energetics of under-water swimming in Adélie penguins (Pygoscelis adeliae). Journal of Comparative Physiology B 161, 285–291 (1991).

21. Beaulieu, M. et al. Can a handicapped parent rely on its partner? An experimental study within Adélie penguin pairs. Animal Behaviour 78, 313–320 (2009).

22. Ropert-Coudert, Y., Wilson, R., Yoda, K. & Kato, A. Assessing performance constraints in penguins with externally-attached devices. Marine Ecology Progress Series 333, 281–289 (2007).

23. Vandenabeele, S. P., Wilson, R. & Grogan, A. Tags on seabirds; how seriously are we considering instrument-induced behaviors? Animal Welfare 20, 559–571 (2011).

24. Ropert-Coudert, Y. et al. Impact of externally attached loggers on the diving behaviour of the king penguin. Physiological and Biochemical Zoology 73, 438–444 (2000).

25. Wilmers, C. C. et al. The golden age of bio-logging: how animal-borne sensors are advancing the frontiers of ecology. Ecology 96, 1741–1753 (2015).

26. CCAMLR. Ecosystem Monitoring Program Standard Methods. (2014).

27. Kenward, R. E. A Manual for Wildlife Radio Tagging. The Auk 118, 812–815 (2001).

28. Wilson, R. P. & McMahon, C. R. Measuring devices on wild animals: what constitutes acceptable practice? Frontiers in Ecology and Evolution 4, 147–154 (2006).

29. Casper, R. M. Guidelines for the instrumentation of wild birds and mammals. Animal Behaviour 78, 1477–1483 (2009).

30. Ratcliffe, N. et al. A leg-band for mounting geolocator tags on penguins. Marine Ornithology 42, 23–26 (2014).

31. Williams, H. J. et al. Optimizing the use of biologgers for movement ecology research. Journal of Animal Ecology 1365-2656.13094 (2019). doi:10.1111/1365-2656.13094

32. Russel, W. M. S. & Burch, R. L. The principles of humane experimental technique. (London: Methuen & Co. Ltd., 1959).

33. Wanless, R. M. & Oatley, T. B. Sixth review of data held by the central data bank for antarctic bird banding, july 1987 – june 1996. Bird Banding (2000).

34. Jackson, S. & Wilson, R. P. The potential costs of flipper-bands to penguins. Functional Ecology 16, 141–148 (2002).

35. Gauthier–Clerc, M. et al. Long–term effects of flipper bands on penguins. Proceedings of the Royal Society of London. Series B: Biological Sciences 271, 423–426 (2004).

36. Saraux, C. et al. Reliability of flipper-banded penguins as indicators of climate change. Nature 469, 203–206 (2011).

37. Dugger, K. M., Ballard, G., Ainley, D. G. & Barton, K. J. Effects of flipper bands on foraging behavior and survival of adélie penguins (Pygoscelis adeliae). The Auk 123, 858 (2006).

38. Le Maho, Y. et al. An ethical issue in biodiversity science: The monitoring of penguins with flipper bands. Comptes Rendus - Biologies 334, 378–384 (2011).

39. Wilson, R. P., Grant, W. S. & Duffy, D. C. Recording devices on free-ranging marine animals: does measurement affect foraging performance? Ecology 67, 1091–1093 (1986).

40. Wienecke, B., Raymond, B. & Robertson, G. Maiden journey of fledgling emperor penguins from the Mawson Coast, East Antarctica. Marine Ecology Progress Series 410, 269–282 (2010).

41. Thiebot, J.-B., Lescroël, A., Barbraud, C. & Bost, C.-A. Three-dimensional use of marine habitats by juvenile emperor penguins Aptenodytes forsteri during post-natal dispersal. Antarctic Science 25, 536–544 (2013).

42. Kooyman, G. L., McDonald, B. I. & Goetz, K. T. Why do satellite transmitters on emperor penguins stop transmitting? Animal Biotelemetry 3, 54 (2015).

43. Goetz, K., McDonald, B. & Kooyman, G. Habitat preference and dive behavior of nonlZlbreeding emperor penguins in the eastern Ross Sea, Antarctica. Marine Ecology Progress Series 593, 155–171 (2018).

44. Labrousse, S. et al. First odyssey beneath the sea ice of juvenile emperor penguins in East Antarctica. Marine Ecology Progress Series 609, 1–16 (2019).

45. Kooyman, G. L. & Ponganis, P. The icing of external recorders during the polar winter. Memoirs of National Institute of Polar Research. Special issue 58, 183–187 (2004).

46. Hays, G. C. et al. Translating marine animal tracking data into conservation policy and management. Trends in Ecology & Evolution 34, 459–473 (2019).

47. Hindell, M. A. et al. Tracking of marine predators to protect Southern Ocean ecosystems. Nature 580, 87–92 (2020).

48. Block, B. Physiological ecology in the 21st century: advancements in biologging science. Integrative and Comparative Biology 45, 305–320 (2005).

49. Costa, D. P., Breed, G. A. & Robinson, P. W. migrations: implications for ecology and conservation. Annual Review of Ecology, Evolution, and Systematics 43, 73–96 (2012).

50. Cooke, S. Biotelemetry and biologging in endangered species research and animal conservation: relevance to regional, national, and IUCN Red List threat assessments. Endangered Species Research 4, 165–185 (2008).

51. Small, C. & Taylor, F. Analysis of albatross and petrel distribution within the CCAMLR Convention Area: Results from the global Procellariiform tracking database. CCAMLR Science 13, 143–174 (2006).

52. CCAMLR. Report of the Twentieth Meeting of the Scientific Committee. (2009).

53. Hays, G. C. et al. Key questions in marine megafauna movement ecology. Trends in Ecology & Evolution 31, 463–475 (2016).

54. Thiebot, J.-B. et al. Adélie penguins’ extensive seasonal migration supports dynamic Marine Protected Area planning in Antarctica. Marine Policy 109, 103692 (2019).

55. Takahashi, A. et al. Migratory movements and winter diving activity of Adélie penguins in East Antarctica. Marine Ecology Progress Series 589, 227–239 (2018).

56. Robertson, G. G. Some field techniques for ecological research on emperor penguins. Marine Ornithology 19, 91–101 (1991).

57. Zimmer, I. Constraints on Foraging and their Consequences for Emperor Penguins PhD. (2007).

58. Cockrem, J. F., Potter, M. A., Barrett, D. P. & Candy, E. J. Corticosterone responses to capture and restraint in emperor and Adélie penguins in Antarctica. Zoological Science 25, 291–298 (2008).

59. Kooyman, G. L. et al. Heart rates and swim speeds of emperor penguins diving under sea ice. The Journal of experimental biology 165, 161–180 (1992).

60. Williams, C. L., Meir, J. U. & Ponganis, P. J. What triggers the aerobic dive limit? Patterns of muscle oxygen depletion during dives of emperor penguins. Journal of Experimental Biology 214, 1802–1812 (2011).

61. Ponganis, P. J., Van Dam, R. P., Marshall, G., Knower, T. & Levenson, D. H. Sub-ice foraging behavior of emperor penguins. Journal of Experimental Biology 203, 3275–3278 (2000).

62. Zimmer, I. et al. Foraging movements of emperor penguins at Pointe Géologie, Antarctica. Polar Biology 31, 229–243 (2007).

63. Zimmer, I. et al. Dive efficiency versus depth in foraging emperor penguins. Aquatic Biology 8, 269–277 (2010).

64. Watanabe, S., Sato, K. & Ponganis, P. J. Activity time budget during foraging trips of emperor penguins. PLoS ONE 7, e50357 (2012).

65. Kirkwood, R. & Robertson, G. The foraging ecology of female Emperor penguins in winter. Ecological Monographs 67, 155–176 (1997).

66. Rodary, D., Bonneau, W., Le Maho, Y. & Bost, C. A. Benthic diving in male emperor penguins Aptenodytes forsteri foraging in winter. Marine Ecology Progress Series 207, 171–181 (2000).

67. Prévost, J. Ecologie du manchot empereur. in Expéditions polaires francaises, (ed. Hermann Press) 204 (1961).

68. Fretwell, P. T. et al. An emperor penguin population estimate: the first global, synoptic survey of a species from space. PLoS ONE 7, e33751 (2012).

69. Ancel, A. et al. Looking for new emperor penguin colonies? Filling the gaps. Global Ecology and Conservation 9, 171–179 (2017).

70. Orians, G. . & Pearson, N. . On the theory of central place foraging. in Analysis of ecological systems (eds. Horn, D., Staris, G. & Mitchell, R.) 155–177 (Columbus: Ohio State University Press, 1979).

71. Stonehouse, B. The Emperor Penguin breeding behaviour and development - FIDS Scientific Report. (1953).

72. Mougin, J. . & Van Beveren, M. Structure et dynamique de la population de Manchots empereurs de la colonie de l’archipel de Pointe Géologie, Terre Adélie. (1979).

73. Kooyman, G. L., Hunke, E. C., Ackley, S. F., Van Dam, R. P. & Robertson, G. Moult of the emperor penguin: Travel, location, and habitat selection. Marine Ecology Progress Series 204, 269–277 (2000).

74. Kooyman, G. L., Siniff, D. B., Stirling, I. & Bengtson, J. L. Moult habitat, pre- and post-moult diet and post-moult travel of Ross Sea emperor penguins. Marine Ecology Progress Series 267, 281–290 (2004).

75. Wienecke, B., Kirkwood, R. & Robertson, G. Pre-moult foraging trips and moult locations of Emperor penguins at the Mawson Coast. Polar Biology 27, 83–91 (2004).

76. Bannasch, R., Wilson, R. P. & Culik, B. Hydrodynamic aspects of design and attachment of a back-mounted device in penguins. The Journal of Experimental Biology 194, 83–96 (1994).

77. Wilson, R. P. et al. Long-term attachment of transmitting and recording devices to penguins and other seabirds. Wildlife Society Bulletin 25, 101–106 (1997).

78. Thiebot, J.-B., Lescroël, A., Pinaud, D., Trathan, P. N. & Bost, C.-A. Larger foraging range but similar habitat selection in non-breeding versus breeding sub-Antarctic penguins. Antarctic Science 23, 117–126 (2011).

79. Le Maho, Y., Delclitte, P. & Chatonnet, J. Thermoregulation in fasting emperor penguins under natural conditions. American Journal of Physiology-Legacy Content 231, 913–922 (1976).

80. Jouventin, P., Barbraud, C. & Rubin, M. Adoption in the emperor penguin, Aptenodytes forsteri. Animal Behaviour 50, 1023–1029 (1995).

81. ATS. General guidelines for visitors to the Antarctic (Attachment to Resolution 3). (2020).

82. Wilson, R. P. & Wilson, M. T. J. Tape: a package-attachment technique for penguins. Wildlife Society Bulletin 17, 77–79 (1989).

83. Ropert-Coudert, Y., Kato, A., Naito, Y. & Cannell, B. L. Individual diving strategies in the little penguin. Waterbirds: The International Journal of Waterbird Biology 26, 403–408 (2003).

84. Chiaradia, A., Ropert-Coudert, Y., Kato, A., Mattern, T. & Yorke, J. Diving behaviour of Little Penguins from four colonies across their whole distribution range: bathymetry affecting diving effort and fledging success. Marine Biology 151, 1535–1542 (2007).

85. Pichegru, L. et al. Diving patterns of female macaroni penguins breeding on Marion Island, South Africa. Polar Biology 34, 945–954 (2011).

86. Le Vaillant, M. et al. How age and sex drive the foraging behaviour in the king penguin. Marine Biology 160, 1147–1156 (2013).

87. Poupart, T. A. et al. Variability in the foraging range of Eudyptula minor across breeding sites in central New Zealand. New Zealand Journal of Zoology 44, 225–244 (2017).

88. Wilson, R. P. et al. Southern rockhopper penguin Eudyptes chrysocome chrysocome foraging at Possession Island. Polar Biology 17, 323–329 (1997).

89. Pütz, K. et al. Post-fledging dispersal of king penguins (Aptenodytes patagonicus) from two breeding sites in the South Atlantic. PLoS ONE 9, e97164 (2014).

90. Williams, T. D. The Penguins: Spheniscidae. The penguins. Bird families of the world. (Oxford University Press, 1995). doi:10.2307/1521878

91. Wienecke, B. & Robertson, G. Foraging space of emperor penguins Aptenodytes forsteri in Antarctic shelf waters in winter. Marine Ecology Progress Series 159, 249–263 (1997).

92. Labrousse, S. et al. Dynamic fine-scale sea icescape shapes adult emperor penguin foraging habitat in east Antarctica. Geophysical Research Letters 46, 11206–11218 (2019).

93. Kooyman, G. L. & Ponganis, P. J. The initial journey of juvenile emperor penguins. Aquatic Conservation: Marine and Freshwater Ecosystems 17, S37–S43 (2007).

94. Trathan, P. et al. Linear tracks and restricted temperature ranges characterise penguin foraging pathways. Marine Ecology Progress Series 370, 285–294 (2008).

95. Sato, K., Shiomi, K., Marshall, G., Kooyman, G. L. & Ponganis, P. J. Stroke rates and diving air volumes of emperor penguins: implications for dive performance. Journal of Experimental Biology 214, 2854–2863 (2011).

96. Wikipedia contributors. Neumayer-Station III. (2021). Available at: https://en.wikipedia.org/w/index.php?title=Neumayer-Station_III&oldid=1024176365. (Accessed: 25th May 2021)

97. McCafferty, D. J., Currie, J. & Sparling, C. E. The effect of instrument attachment on the surface temperature of juvenile grey seals (Halichoerus grypus) as measured by infrared thermography. Deep Sea Research Part II: Topical Studies in Oceanography 54, 424–436 (2007).

98. Gilbert, C., Robertson, G., Le Maho, Y., Naito, Y. & Ancel, A. Huddling behavior in emperor penguins: Dynamics of huddling. Physiology and Behavior 88, 479–488 (2006).

99. Walker, K. A., Trites, A. W., Haulena, M. & Weary, D. M. A review of the effects of different marking and tagging techniques on marine mammals. Wildlife Research 39, 15 (2012).

100. Wanless, S., Frederiksen, M., Daunt, F., Scott, B. E. & Harris, M. P. Black-legged kittiwakes as indicators of environmental change in the North Sea: Evidence from long-term studies. Progress in Oceanography 72, 30–38 (2007).

101. Orgeret, F., Weimerskirch, H. & Bost, C.-A. Early diving behaviour in juvenile penguins: improvement or selection processes. Biology Letters 12, 20160490 (2016).

102. Enstipp, M. R. et al. Apparent changes in body insulation of juvenile king penguins suggest an energetic challenge during their early life at sea. The Journal of Experimental Biology 220, 2666–2678 (2017).

103. Thiebot, J.-B. & Pinaud, D. Quantitative method to estimate species habitat use from light-based geolocation data. Endangered Species Research 10, 341–353 (2010).

104. Ballard, G. et al. Responding to climate change: Adélie Penguins confront astronomical and ocean boundaries. Ecology 91, 2056–2069 (2010).

105. Dunn, M. J., Silk, J. R. D. & Trathan, P. N. Post-breeding dispersal of Adélie penguins (Pygoscelis adeliae) nesting at Signy Island, South Orkney Islands. Polar Biology 34, 205–214 (2011).

106. Pierce, A. J., Stevens, D. K., Mulder, R. & Salewski, V. Plastic colour rings and the incidence of leg injury in flycatchers (Muscicapidae, Monarchidae). Ringing and Migration 23, 205–210 (2007).

107. Griesser, M. et al. Causes of ring-related leg injuries in birds - evidence and recommendations from four field studies. PLoS ONE 7, (2012).

108. Costantini, D. & Møller, A. P. A meta-analysis of the effects of geolocator application on birds. Current Zoology 59, 697–706 (2013).

109. Richter, S. et al. A remote-controlled observatory for behavioural and ecological research: A case study on emperor penguins. Methods in Ecology and Evolution 9, 1168–1178 (2018).

110. Gendner, J.-P. et al. Automatic weighing and identification of breeding penguins. In 4th European international conference on wildlife telemetry, llis Horwood Ltd 29:30 (1992).

111. Gendner, J.-P., Gauthier-Clerc, M., Le Bohec, C., Descamps, S. & Le Maho, Y. A new application for transponders in studying penguins. Journal of Field Ornithology 76, 138–142 (2005).

112. Le Bohec, C. et al. Population dynamics in a long-lived seabird: I. Impact of breeding activity on survival and breeding probability in unbanded king penguins. Journal of Animal Ecology 76, 1149–1160 (2007).

113. Cristofari, R. et al. Climate-driven range shifts of the king penguin in a fragmented ecosystem. Nature Climate Change 8, 245–251 (2018).

114. Thiebot, J.-B., Cherel, Y., Trathan, P. N. & Bost, C. A. Inter-population segregation in the wintering areas of macaroni penguins. Marine Ecology Progress Series 421, 279–290 (2011).

115. Thiebot, J.-B. et al. Mates but not sexes differ in migratory niche in a monogamous penguin species. Biology Letters 11, 1–4 (2015).

116. Enstipp, M. R. et al. The dive performance of immature king penguins following their annual molt suggests physiological constraints. The Journal of Experimental Biology 222, jeb208900 (2019).

117. Forin-Wiart, M.-A., Enstipp, M. R., Le Maho, Y. & Handrich, Y. Why implantation of bio-loggers may improve our understanding of how animals cope within their natural environment. Integrative Zoology 14, 48–64 (2019).

